# In vitro To In vivo : Bidirectional and High-Precision Generation of In Vitro and In Vivo Neuronal Spike Data

**DOI:** 10.1101/2025.03.26.645366

**Authors:** Masanori Shimono

## Abstract

Spontaneous neural activity is often dismissed as noise, yet predicting its future evolution remains difficult, especially across recording preparations. We study transfer between in vitro and in vivo multineuronal spike trains with a Transformer that treats spikes as sparse point processes and does not require cell-to-cell correspondence across datasets. Because binned spike trains are extremely sparse and imbalanced, likelihood-based objectives can be dominated by the majority no-spike class under domain shift. Introducing a Dice objective that maximizes spike-event overlap stabilizes training and substantially improves performance, including in cross-domain prediction and generation. Training on one domain and evaluating on another yields a quantitative predictability score and a predictability matrix. We show that (i) a performance-based embedding organizes datasets by transferability; (ii) only specific in vitro and in vivo pairs support reliable cross-domain generation, whereas mismatched pairs degrade despite strong within-domain accuracy; (iii) attention and gradient-based importance analyses reveal how the model shifts emphasis across neurons and time; and (iv) the predictability matrix guides source–target selection for generating diverse activity patterns. This framework supports systematic tests of transferability and may help prioritize datasets for translational studies.

## 1. Introduction

The nervous system transmits signals and performs computation through the spiking activity of neurons. To understand how information is encoded at the level of large populations, it is essential to characterize the spatiotemporal “recipes” that emerge from interactions among many neurons. Even without explicit cognitive tasks, the brain exhibits spontaneous activity. Across development, accumulated experience and internal constraints shape this ongoing activity, embedding a repertoire of latent patterns that can be rapidly recruited when external inputs arrive [1,2]. Yet precisely because spontaneous activity is rich and not time-locked to an external event, it has often been treated as “noise,” and its moment-to-moment evolution remains challenging to predict and compare across settings [2,3]. In this sense, spontaneous activity may carry a compressed trace of an individual’s past experience, while remaining difficult to model because of its complexity.

Population dynamics are not arbitrary: they are constrained by anatomical wiring and mesoscale organization. Large-scale anatomical and functional connectomics has sharpened the link—close but nontrivial—between structure and activity in cortex [4–7]. Complementary network-level analyses quantify how structure–function coupling varies across space and time, reinforcing the view that spontaneous dynamics are shaped by an anatomical scaffold [6]. Moreover, effective interactions can sometimes be inferred from activity time series, implying that partially predictable dynamical motifs may be embedded in spontaneous fluctuations [8,9].

From a translational viewpoint, a central challenge is whether dynamical regularities learned in one experimental preparation transfer to another. Crossing preparations (for example, from in vitro slices to in vivo recordings) becomes especially difficult because brain state changes, nonstationarity, extreme sparsity of binned spikes, and the lack of neuron-to-neuron correspondence can all degrade generalization. Despite the complementary strengths of in vitro controllability and in vivo physiological realism, connecting the two quantitatively is still not seamless in practice, and guidance for coordinated experimental design remains limited [10,11].

Methodologically, sequence models have begun to capture population spiking at scale. We previously demonstrated that LSTM-based models can generate spontaneous activity across in vitro cortical regions, supporting the idea that transferable motifs exist even without explicit stimuli [12]. More recently, Transformer architectures and neural-data variants (e.g., NDT, STNDT, NDT2) have enabled efficient parallel modeling of binned spike trains [13–16]. However, dominant evaluations still emphasize within-domain reconstruction or decoding, leaving time-resolved, cross-preparation transfer insufficiently quantified.

Here, we present a Transformer-based framework for bidirectional domain transfer between in vitro and in vivo spontaneous spike trains. To address severe class imbalance, we optimize a Dice-based objective [17]. We introduce a time-resolved transfer evaluation that emphasizes low false-positive regimes, construct a predictability matrix and low-dimensional embeddings to summarize cross-dataset similarity, and provide model-based analyses that connect transferable predictability to circuit-level constraints.

## 2. Materials and Methods

### 2.1. Data Acquisition

#### 2.1.1. In Vitro Data

The in vitro data were collected from cortical slices of C57BL/6 mice (3–5 weeks old) using high-density microelectrode arrays (HD-MEAs, MaxWell Biosystems Co.), following the methodologies described in Nakajima et al. (2023) and Matsuda et al. (2023).

Prior to tissue extraction, each mouse was deeply anaesthetized with 1–1.5 % isoflurane delivered via inhalation. While full surgical anesthesia was confirmed (no pedal reflex), euthanasia was performed by cervical dislocation. The brain was then rapidly removed and immersed in ice-cold cutting solution for slice preparation.

From the recorded electrical signals, spike sorting was carefully conducted using Spyking Circus software, allowing for the extraction of neural activity data from approximately 1,000 adjacent cells. To ensure the accuracy of cortical region identification, these electrophysiological recordings were further aligned with immunostained images, from which 128 representative neurons were selected for analysis.

In previous studies, cortical data were categorized into 16 groups (eight per hemisphere). However, considering that in vivo data were exclusively recorded from the left hemisphere, we refined our dataset selection accordingly. To maintain consistency and data quality, we limited our analysis to six groups of in vitro data from the left hemisphere (Table 1 and Figure 1).

**Figure 1.**
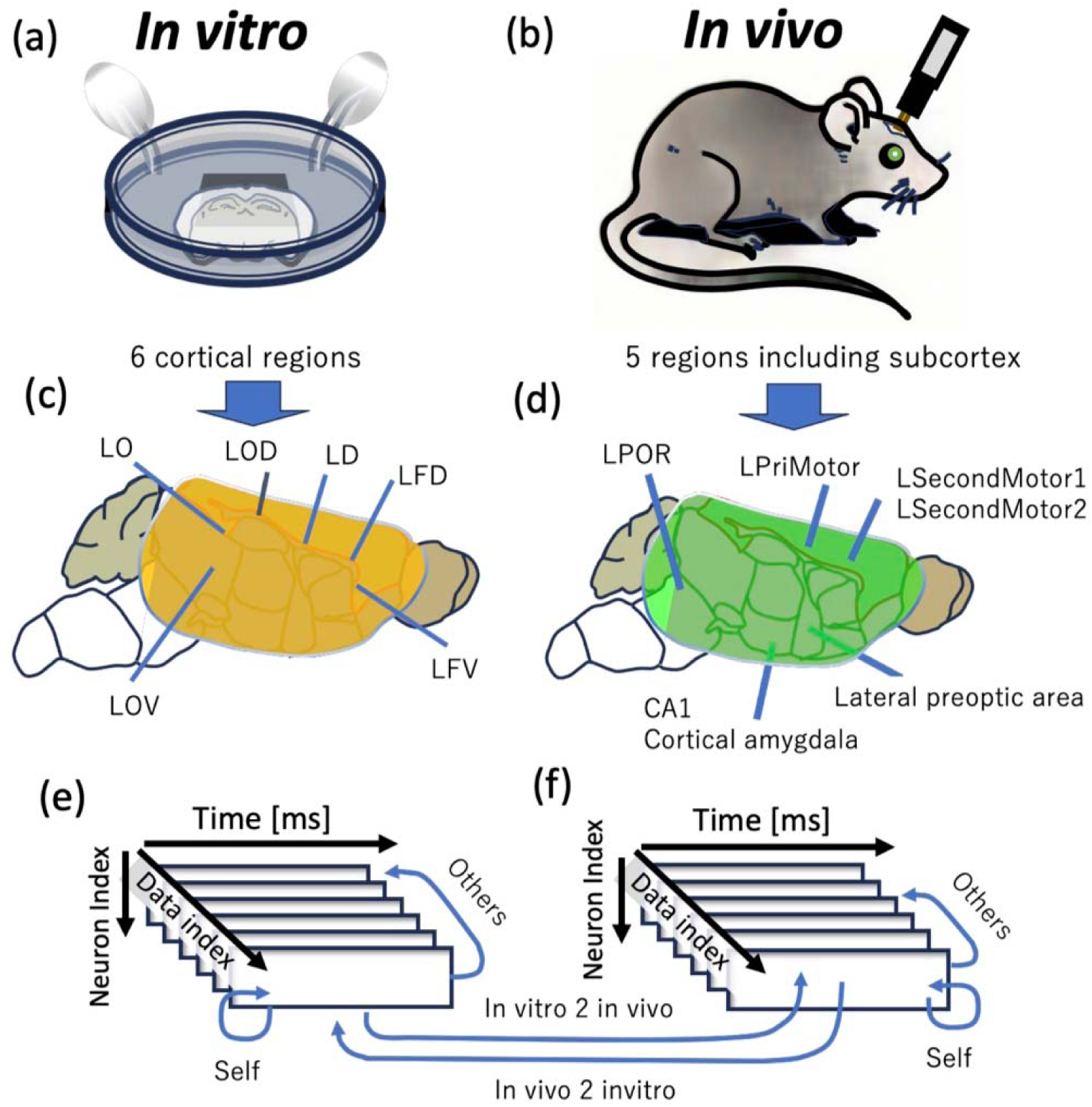
Illustration of the Data Sets Used in This Study. Figure 1 provides an overview of the data sets utilized in this study. Panel (a) depicts the setup of an in vitro electrophysiological experiment. A brain acute slice is placed on an electrode, perfused with artificial cerebrospinal fluid (ACSF), and continuously bubbled with oxygen-enriched gas while neural activity is recorded. Panel (b) illustrates the setup for an in vivo electrophysiological experiment. In this case, electrodes are inserted into targeted brain regions of a living mouse to measure neural activity. Panels (c) and (d) indicate the brain regions recorded during the in vitro and in vivo experiments, respectively. These panels display abbreviated names representing each recorded brain region. The meanings of these abbreviations and the specific brain regions they correspond to are summarized in Table 1. Panels (e) and (f) represent two-dimensional neural activity data obtained from each brain region over time, with individual squares corresponding to neuronal activity at different time points. The blue arrows illustrate the generated pattern pairs within each experiment. Within both (e) and (f), the blue arrows indicate self-generation from the source data, as well as generation towards other regions (self-to-self and self-to-others). Additionally, inter-experimental generation pathways between in vitro and in vivo data are depicted, specifically in vitro-to-in vivo and in vivo-to-in vitro generation, as shown by the interconnecting blue arrows between panels (e) and (f). The recorded brain regions and abbreviations used in this study are summarized in Table 1.

**Table 1.**
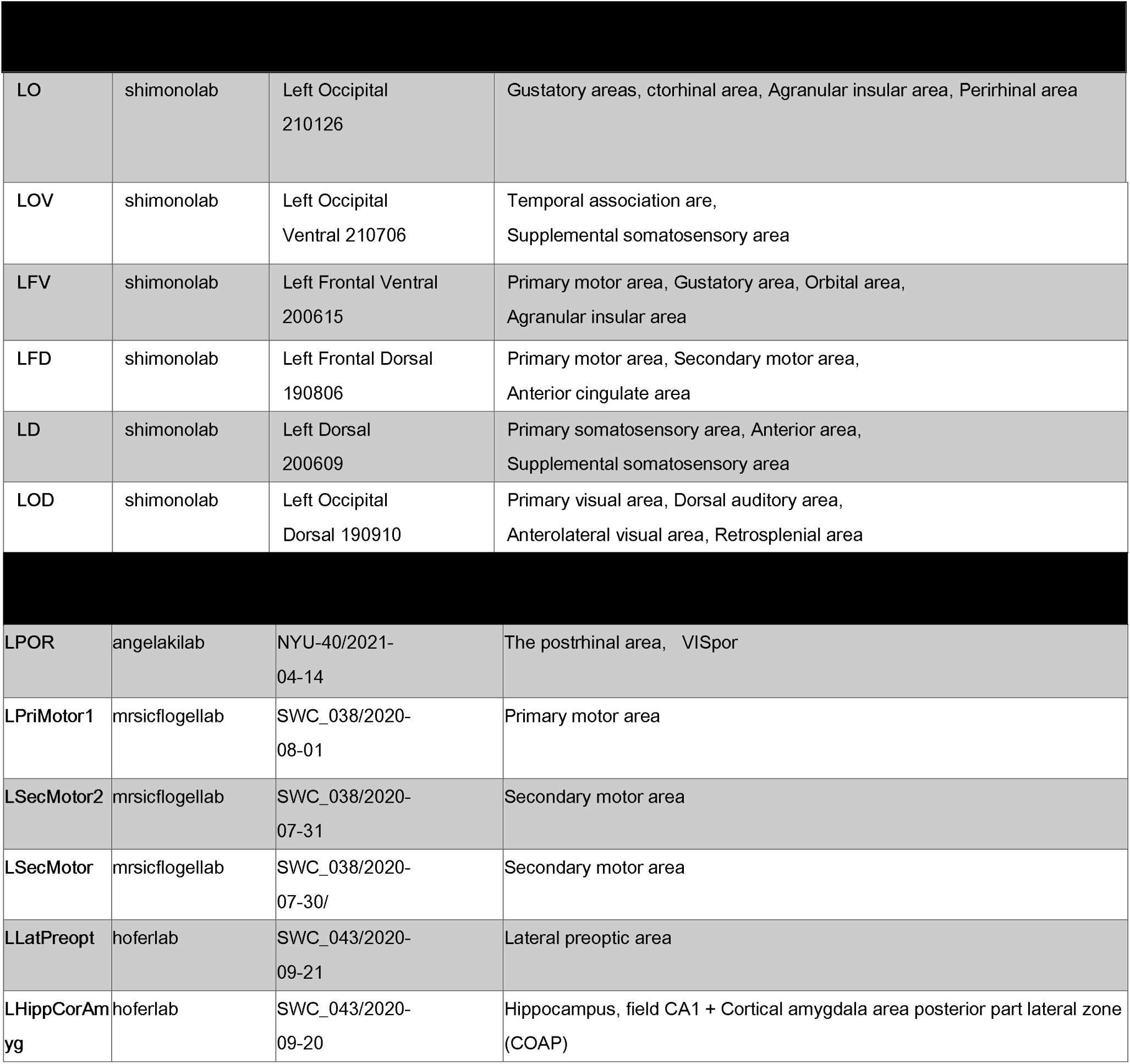
List of Brain Regions Corresponding to Experimental Data. Table 1 presents a comprehensive list of brain regions corresponding to the in vitro and in vivo experimental data. From left to right, the table includes: (1) the dataset name as used in this study, (2) the laboratory responsible for data acquisition, (3) supplementary dataset information such as measurement dates, and (4) the specific brain regions recorded. The in vitro data were obtained by the Shimono Lab, with recordings conducted from multiple regions of the left cerebral hemisphere. In contrast, the in vivo data were collected by the Angelaki Lab, Mrsic-Flogel Lab, and Hofer Lab, covering a wide range of brain regions, including the motor cortex, visual cortex, hippocampus, and amygdala. For further reference, see: https://viz.internationalbrainlab.org/app?spikesorting=ss_2024-05-06&dset=bwm&pid=9915765e-ff15-4371-b1aa-c2ef1db8a254&tid=5&cid=388&qc=1.

To minimize the impact of non-stationarity, we excluded the first 30 minutes of recordings from each in vitro session. The subsequent 7.5 minutes were then segmented into training and test datasets. A detailed list of the in vitro datasets used in this study is provided in Table 1. For further details regarding the dataset, refer to the cited studies [Nakajima et al., 2023; Matsuda et al., 2023].

#### 2.1.2. In Vivo Data

The in vivo data were obtained from the International Brain Laboratory (IBL), specifically using passive stimulation protocol recordings from C57BL/6 mice (15–63 weeks old) [International Brain Laboratory et al., 2023; 2025]. The in vivo data used in this study were recorded between 111 to 442 days of age (mean: 34.43 weeks, median: 26.0 weeks), based on data collected before 2022.For analysis, we focused on 7.5 minutes of spontaneous activity recorded at the beginning of each session.

The International Brain Laboratory compiles data from multiple laboratories worldwide. In this study, we selected datasets containing spontaneous activity recorded in several of these laboratories for analysis. The full list of datasets used is summarized in Table 1.

All electrophysiological recordings in this dataset were collected using Neuropixels 1.0 multi-electrode probes [Jun et al., 2017]. Spike sorting was performed using a motion-corrected three-dimensional spike localization method optimized for this electrode type [Boussard et al., 2021]. Further details on experimental procedures and data processing pipelines for extracting neural activity time series can be found in the IBL experiment documentation: https://int-brain-lab.github.io/iblenv/notebooks_external/loading_passive_data.html

### 2.2. Analysis Methods

#### 2.2.1. Data Segmentation

In both in vitro and in vivo datasets (Figure 2a,b), we extracted 7.5-minute segments of spike activity recorded from 128 neurons to predict neural activity patterns within specific brain regions.

**Figure 2.**
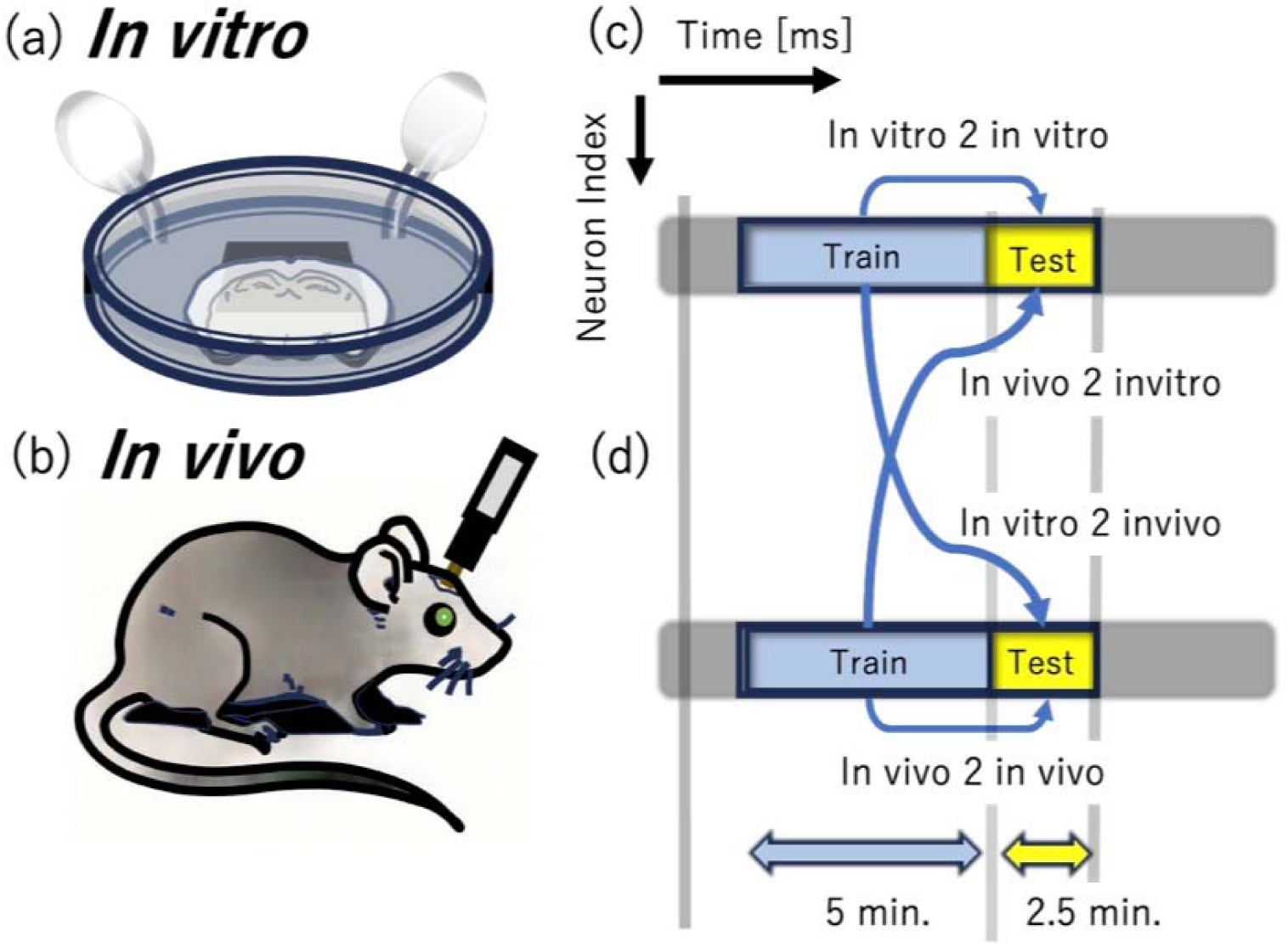
This figure explains the time duration of extracted data and the method of dividing data into training and test sets. (a) and (b) show in vitro data in the upper panels and in vivo data in the lower panels. For both (c) in vitro and (d) in vivo data, we extract 5-minute segments for training data and 2.5-minute segments for test data to conduct learning and prediction. The combinations of choosing either in vitro or in vivo data for training and testing are classified as in vitro2in vitro, in vivo2in vivo, in vitro2in vivo, and in vivo2in vitro.

Each 7.5-minute dataset was divided into a 5-minute training set and a 2.5-minute test set (Figure 2c). This segmentation was a crucial step for assessing the model’s generalization ability—after training the model on the training set, its predictive performance was evaluated using the test set.

For within-domain generation (e.g., in vitro to in vitro or in vivo to in vivo), the training and test data were drawn from a continuous 7.5-minute session. However, for cross-domain generation (e.g., in vitro to in vivo), the model was trained and tested on entirely different 5-minute and 2.5-minute segments, ensuring no overlap between training and test datasets.

This segmentation into 5 minutes for training and 2.5 minutes for testing, ensuring no overlap or double dipping in model evaluation. Our framework does not employ iterative re-training on generated outputs—not due to a fear of model collapse, but because faithful reproduction of original data characteristics is central to enabling direct, dynamics-preserving comparisons across datasets.

#### 2.2.2. Transformer Model and Loss Function Selection

The primary analytical algorithm employs a single-layer Transformer-based model [Vaswani et al., 2017]. The Transformer model implements multi-head attention with 32 heads and accepts 5,000 input tokens. During training, we also applied a dropout rate of 10% to the connections. Although we tested multi-layer Transformer models prior to final evaluation, these showed slower receiver operating characteristic (ROC) curve growth and generally poorer performance, leading us to ultimately adopt a single-layer Transformer model. During training, we utilized the Dice loss function to maximize prediction accuracy for binary (0 and 1) data. This loss function is particularly suitable for achieving stable learning in the presence of class imbalance. We adopted this loss function as it performed better than the focal-loss baseline tested here, and showed favorable behavior in preliminary comparisons.

#### 2.2.3. Attention Map

To gain deeper insights into the information learned by the Transformer model during prediction generation, we quantitatively evaluated the model’s learned information and predictive features by combining attention map dynamics with gradient-based importance. Here, we explain the relevant aspects of the Transformer model’s internal structure.

The self-attention mechanism, which achieved breakthrough results particularly in Natural Language Processing (NLP) [Vaswani et al., 2017], is considered the central mechanism in Transformer model learning. At its core is the attention map, which represents a matrix indicating how much attention each input element (token) should pay to other input elements. The attention map is mathematically expressed as:

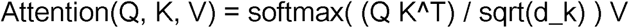

where Q = X W^Q, K = X W^K, V = X W^V, d_k is the key dimension, and S = (Q K^T)/sqrt(d_k) gives the similarity scores.

where Q (Query) and K (Key) are linearly transformed representations of the input sequence X. The (i, j) element of the similarity matrix S, denoted S_{i,j}, represents the similarity between token i and token j; a higher value indicates a stronger influence of input token i on the processing of token j.

The similarity scores S are scaled by sqrt(d_k) and normalized with softmax to yield the attention weights A. These weights are then used to compute a weighted sum of V to produce the final output.

While attention maps have been considered fundamental to success in various tasks including machine translation [Vaswani et al., 2017], document summarization [Liu & Lapata, 2019], and image recognition [Dosovitskiy et al., 2021], recent studies have indicated that attention maps alone are insufficient to explain transformer learning [Jain, Wallace, 2019; Serrano, Serrano, 2019]. Indeed, our research revealed phenomena that cannot be explained by attention maps alone, leading us to calculate the model’s gradient-based importance.

#### 2.2.4. Gradient-based Importance and Attention-weighted Importance

Gradient-based importance measures the influence of each input dimension on the loss function and is defined as follows: where represents the loss function and is the kth element of the feature when Query pays attention to Key.

In other words, represents the gradient sum over, indicating how much the intermediate feature influences the loss function. This summation allows us to obtain the global importance of each input element.

This method computes global importance for each input element independently of the local attention distribution provided by attention maps, directly reflecting the relationship with the loss function.

To observe token interactions, we integrated this Importance with the Attention Map as follows:

We term this quantity attention-weighted importance and use it for evaluation. Here, indicates relative attention between inputs, and is the transpose of the importance vector calculated based on gradient information. This operation combines the “inter-input attention distribution” shown by the Attention Map with the “importance in the loss function” indicated by gradient information, achieving higher interpretability.

#### 2.2.5. Generation Data Evaluation Method

During the testing phase, we generated predictions by repeatedly predicting 1ms ahead using past test data while keeping the trained model fixed. To evaluate the generated data, we quantified the model’s discriminative ability using the Area Under the Curve (AUC) of the Receiver Operating Characteristic (ROC) curve, which shows the relationship between false positive and true positive rates.

#### 2.2.6. Network Layout Optimization in Three-Dimensional Space

This study mapped brain regions in three-dimensional space based on ROC-AUC values, which represent the ease of mutual generation between regions. Specifically, we treated the inverse of ROC-AUC as distances to create a distance matrix between brain regions, then sought a configuration that minimized overall energy while maintaining these prescribed distances between nodes as much as possible. The optimization process followed these steps: First, initial placement: Nodes were randomly positioned in three-dimensional space. Second, energy function definition: Sum of squared differences between actual and desired distances for all node pairs.

Third, position optimization: Node positions were optimized to minimize this energy function (Figure 4f)

### 2.3. Ethics statement

Ethical approvals and compliance statements for the in vitro and in vivo experiments are provided in the Institutional Review Board Statement (Back matter).

## 3. Results

### 3.1. Evaluation and Comparison of Loss Functions During Training

In this study, we trained our model using in vitro data measured from slices of seven cortical regions in the left hemisphere, as well as six in vivo datasets recorded from either the cortex or hippocampus of the left hemisphere. To optimize the learning process, we employed Dice Loss as the loss function. As a result, across all training datasets, the area under the ROC curve (AUC) reached 0.92 ± 0.06 (Figure 3). Due to severe class imbalance, we report ROC-AUC as the primary metric. During training, the true-positive and true-negative rates approached ∼90% on average, but these should be interpreted as secondary to ROC/PR metrics. This high level of accuracy was difficult to achieve with many alternative methods beyond the use of Dice Loss.

**Figure 3.**
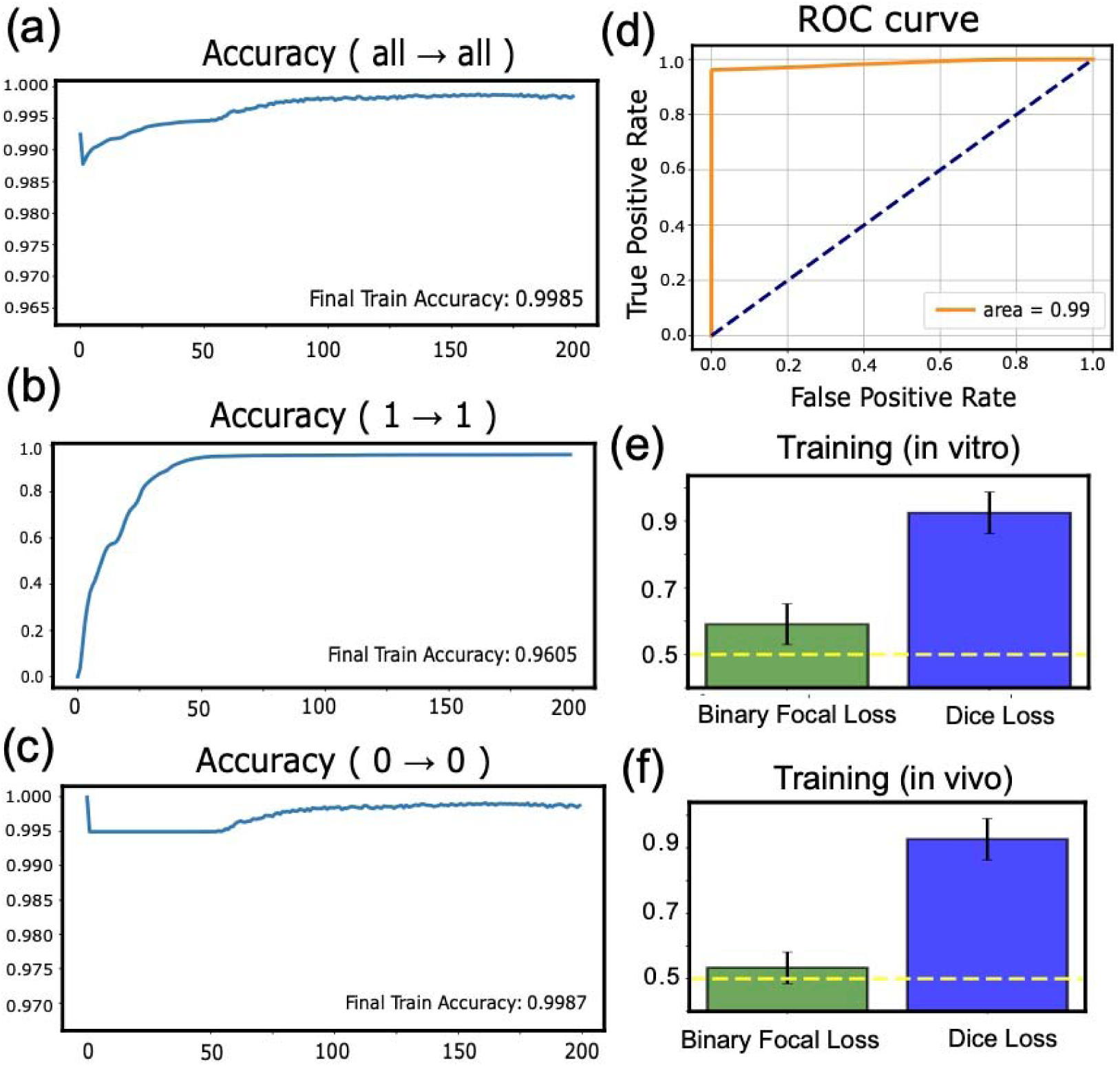
Learning Process and Training Results. Panel (a) illustrates the progression of accuracy during the training process across all datasets over 200 epochs. Panel (b) depicts the changes in accuracy specifically for correctly predicting non-spiking (0) states throughout training, while panel (c) shows the corresponding accuracy changes for spiking (1) states. Panel (d) presents a representative ROC curve after training. The curve initially extends vertically, indicating an increase in true positives, before bending at a later stage. This pattern indicates high discriminative ability in distinguishing spiking events. Panels (e) and (f) compare the training accuracy (ROC-AUC) for in vitro and in vivo data using Binary Focal Loss and Dice Loss, represented as bar graphs. The results clearly indicate that the use of Dice Loss improved prediction accuracy in our datasets, outperforming Binary Focal Loss.

Neural spike data present a unique challenge due to the extremely low frequency of the “1” state (spiking events). Consequently, predicting the occurrence of these essential “1” states is highly difficult. By employing Dice Loss as the error function, we effectively corrected the imbalance between the occurrence frequencies of 0 and 1, demonstrating its significant utility in this context (Figure 3a,b, c).

To further assess the effectiveness of Dice Loss, we compared its performance with the commonly used Binary Focal Cross Entropy Loss (Figure 3). In this study, we quantitatively evaluated predictive performance using the Area Under the Curve (AUC) of the Receiver Operating Characteristic (ROC) curve. With Binary Focal Cross Entropy Loss, statistical significance was obtained only in self-prediction tasks using in vitro data. However, when Dice Loss was applied, the model achieved AUC values exceeding 0.9, demonstrating high prediction accuracy not only for in vitro data but also for in vivo data (Figure 3e,f).

These results indicate that Dice Loss effectively overcomes the challenge posed by the frequency imbalance between 0 and 1, significantly improving learning performance and achieving high-precision training outcomes.

### 3.2. Results of Generation

Our computational experiments demonstrated that training with Dice Loss yielded successful results. Specifically, the training performance achieved high scores of 0.92±0.06 for in vitro data and 0.93±0.06 for in vivo data. The corresponding time series examples are shown in Figures 4c and 4d.

Figures 1e,f, and 3a illustrate progressive learning and strong generalization across datasets. Even in the lowest-scoring case, the ROC-AUC for in vitro and in vivo data generation was 0.70 ± 0.09, as summarized in Table 2 and Figure 4b.

**Table 2.** Performance Comparison Across Training Conditions. This table presents the performance metrics (mean ± standard deviation) under different training conditions using in vitro and in vivo data. Both in vitro and in vivo data showed high performance during training with their respective data (0.92±0.06, 0.93±0.06). Performance tended to decrease when using data from different sources compared to using self-data. However, generation between different environments also achieved performance above chance (AUC>0.5) and above our baseline comparisons. https://viz.internationalbrainlab.org/app?spikesorting=ss_2024-05-06&dset=bwm&pid=9915765e-ff15-4371-b1aa-c2ef1db8a254&tid=5&cid=388&qc=1.

| Conditions | Performance |
| --- | --- |
| In vitro (training) | $0.92\pm0.06$ |
| In vivo (training) | $0.93\pm0.06$ |
| In vitro 2 invitro ( self data ) | $0.93\pm0.05$ |
| In vitro 2 in vitro ( others ) | $0.75\pm0.07$ |
| In vivo 2 in vivo ( self data ) | $0.77\pm0.10$ |
| In vivo 2 in vivo ( others ) | $0.74\pm0.10$ |
| In vitro 2 in vivo | 0.70±0.09 |
| In vivo 2 in vitro | 0.80±0.10 |

To analyze these results more comprehensively, we visualized the ROC-AUC scores for all combinations as a color map (Figure 4e). Figure 4f depicts the ROC-AUC color map matrix as a network diagram, where the inverse of the connection strength is treated as a distance and the positions are optimized in three-dimensional space. From these visualizations (Figure 4e,f), we identified five key findings:

First, in vitro to in vitro generation showed favorable performance with ROC-AUC scores around 0.93 between identical regions. While this exceptionally high performance was unexpected, the relatively strong performance in this combination was anticipated.

**Figure 4.**
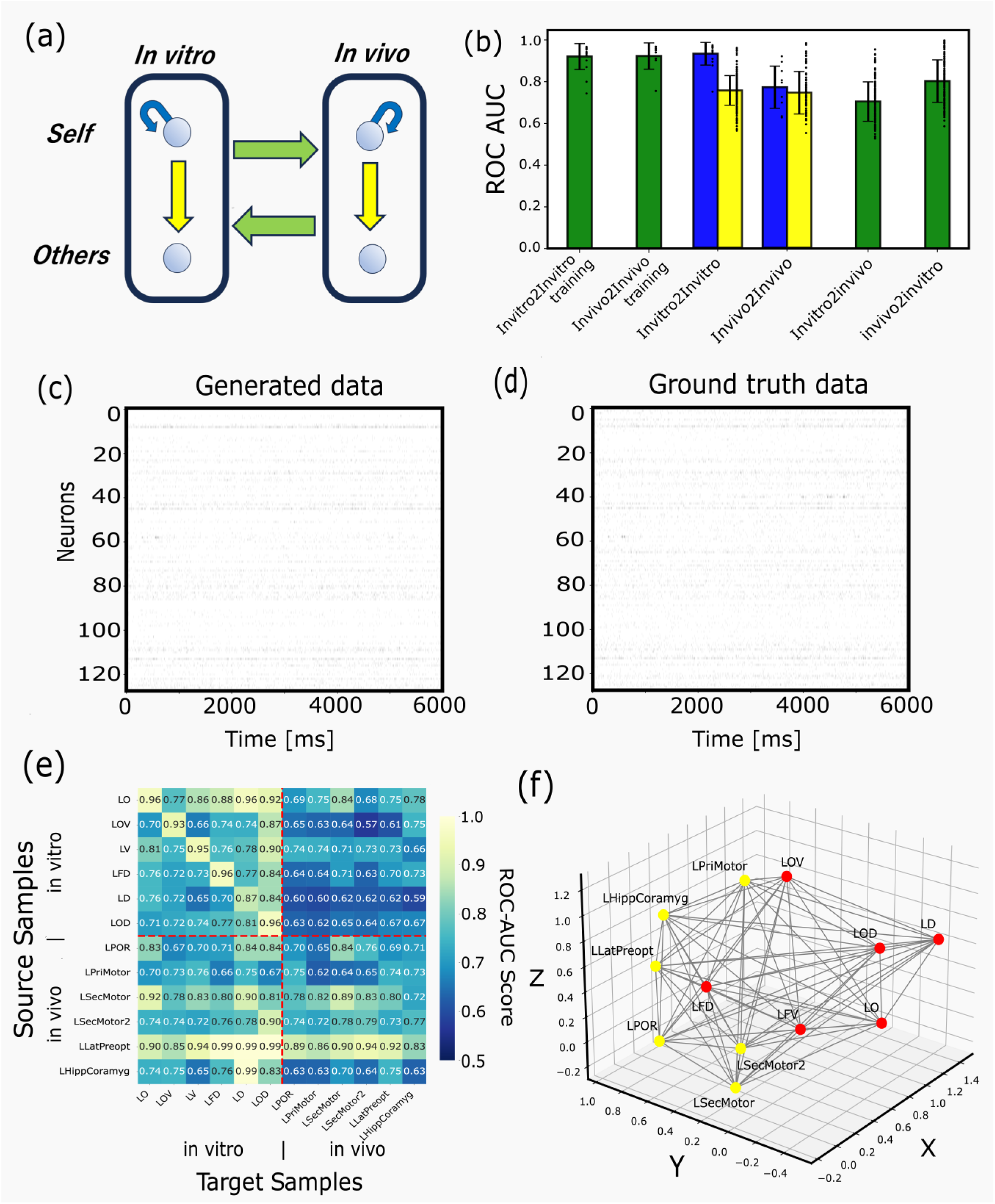
Results of Prediction-Generation Process. Panel (a) presents a schematic diagram summarizing the various combinations of training and prediction data, effectively reframing Figures 1e,f. For in vitro data (circle at top left), both prediction of “Self” (curved blue arrow) between past and future own states and “Others” prediction (downward yellow arrow) to future states of other in vitro data are possible. Similarly, for in vivo data (circle at top right), both “Self” prediction-generation (curved blue arrow) and “Others” prediction-generation (downward yellow arrow) exist. Bidirectional generations between in vitro and in vivo data also exist, termed in vitro2in vivo and in vivo2in vitro for rightward and leftward generation, respectively. Panel (b) displays bar graphs showing performance metrics for both training performance and prediction performances for the various combinations of training and prediction data outlined in (a). The leftmost two bars represent final training performance for in vitro and in vivo data. The next pair shows in vitro2in vitro and in vivo2in vivo generation, with each pair comprising “self” prediction (left bar) and “others” prediction (right bar). The rightmost two bars represent in vitro2in vivo and in vivo2in vitro generation. Panels (c) and (d) present examples comparing generated time series with ground truth, with time on the horizontal axis and neuron index on the vertical axis. Panel (e) displays a two-dimensional representation of performance metrics for combinations of training data (vertical axis) and prediction data (horizontal axis) across 12 datasets (6 in vitro and 6 in vivo). Red dotted lines divide the color map into quadrants: in vitro to in vitro generation (top left), in vitro to in vivo generation (top right), in vivo to in vitro generation (bottom left), and in vivo to in vivo generation (bottom right). The bar graph in (b) represents these grouped results, with ROC-AUC scores on the vertical axis, separated into diagonal and non-diagonal components where applicable. Region abbreviations follow the IDs listed in Table 1. Panel (f) represents the matrix from (e) reconfigured as a network diagram. It maps the data samples by optimizing positions after regarding the inverse of the AUC (used to measure prediction accuracy) as a distance metric in three-dimensional space. The red circular nodes represent in vitro data, while the yellow circular nodes represent in vivo data. The abbreviations for the region names follow the same conventions as in Table 1.

Second, interestingly, in vivo to in vivo generation did not demonstrate particularly superior performance between identical regions. We did not find evidence for greater non-stationarity in the in vivo data based on an Augmented Dickey-Fuller test (p=0.0083); however, this result depends on our preprocessing choices and the time windows analyzed [Dickey & Fuller, 1981]. Our findings suggest that in vivo data exhibits stronger inter-regional influences compared to in vitro data, leading to variations in spike patterns.

Third, in vivo to in vitro generation outperformed in vitro to in vivo generation. This suggests that in vivo data encompasses the activity repertoire of in vitro data while partially acquiring new activity patterns. However, bidirectional comparisons indicate a non-trivial overlap with in vitro data in our setting.

Fourth, the lateral preoptic area (LPO) data showed relatively strong cross-region predictability within our sample. This finding will be extensively discussed in the Discussion section. In contrast to the lateral preoptic area, the cerebellum was less effective as a seed and more readily generated from other data in our sample; this may reflect simpler patterns in these recordings, but broader data would be needed to generalize.

Fifth, we could observe that in vitro data tends to cluster together with other in vitro data. At the same time, regions related to the cortical motor area, regardless of whether they are in vitro or in vivo, are concentrated in the central part. These characteristics support the idea that the ROC-AUC-based mapping meaningfully arranges the diversity of activity. Further insights can be gained by comparing this with Figures 1c and 1-d.

### 3.3. Analysis of Information Learned by the Model

To gain insight into the predictive mechanisms of the Transformer model, we conducted an analysis of the internal processing of information within the model. Here, to maintain focus and avoid unnecessary complexity, we limited our analysis to the case of in vitro to in vivo predictions.

In translation tasks using language models, it is well known that the correspondence between input words (before translation) and output words (after translation) is captured in the Attention Map [Bahdanau et al., 2015; Voita et al., 2019]. Therefore, we began our analysis by examining the Attention Map (see the “Attention Map” subsection in the Methods section). Since our study deals with binary sequences (0s and 1s), we analyzed the relationship between the firing rate of the output signals and the weighted Attention Map. Specifically, to investigate how past information influences predictions, we examined how the average weight changes relative to the diagonal components of the Attention Map, which indicate time shifts from the present moment (Figure 5a). The results revealed a clear peak along the diagonal component, suggesting that the model heavily relies on data from immediately preceding time points for its predictions. However, off-diagonal components were also observed, indicating that the model may be assigning supplementary attention to specific past moments.

**Figure 5.**
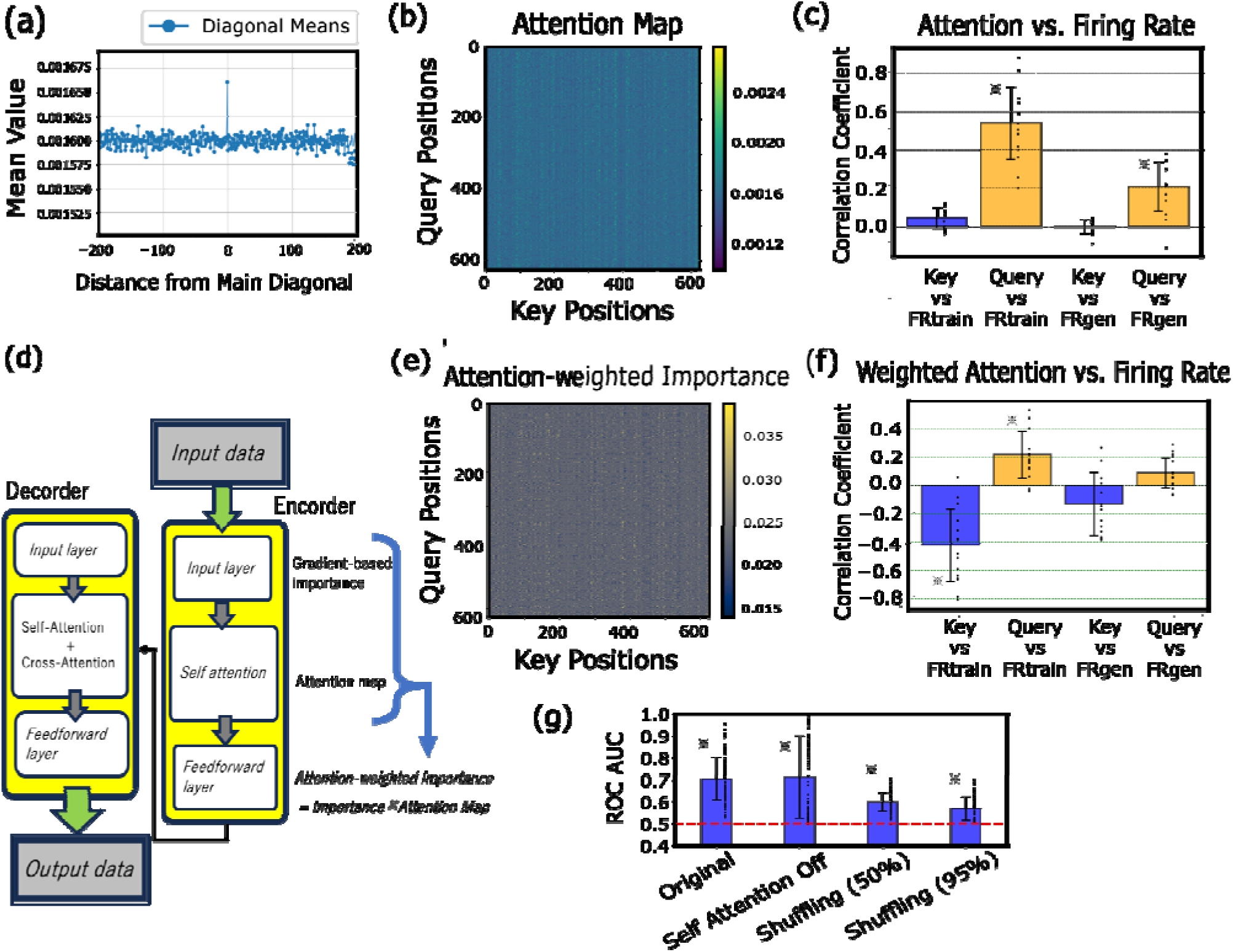
Analysis of Information Learned by the Transformer Model. Panel (a) illustrates representative attention maps within the Transformer model. Panel (b) presents examples of attention-weighted importance maps, which are attention maps weighted by importance scores. Panel (c) shows a schematic diagram depicting the relative relationships between input signals, attention maps, attention-weighted importance, and output signals. Panel (d) demonstrates the intensity distribution of deviations from the diagonal components in the attention maps. A consistent peak near zero was observed across all datasets. Panel (e) displays correlation plots comparing firing rates between training and generated data for both Key and Query sides of the attention maps. Results are shown as means with standard deviations, with individual data points plotted alongside error bars. Asterisks (*) indicate statistical significance at p<0.01 (t-test). Panel (f) presents correlation plots comparing firing rates between training and generated data for both Key and Query sides of the attention-weighted importance maps. The format follows that of panel (e). Panel (g) displays four bar graphs. From left to right, they represent the following conditions: the original in vitro to in vivo generation case (second bar from the right in Figure 4b), a case where the attention mechanism is disabled, and prediction performance when the input data was shuffled across cells within each time slice for either 50% or 95% of the temporal window, extending from future time points to past ones. Asterisks (*) indicate statistical significance at p<0.01 (t-test) when compared against the zero-correlation line.

Next, we compared the Attention Map with the firing rates of input signals used during training. The results showed that the Attention Map correlated significantly with the query-side firing rate of the training data, whereas no significant correlation was found with the key-side firing rate (Figure 5b,c). To assess the critical role of the Attention mechanism in this learning process, we analyzed performance changes when the Attention mechanism was disabled. However, the results indicated that learning performance did not significantly deteriorate (Figure 5g, left two bars).

Given this outcome, we expanded our analysis beyond the Attention Map and introduced an importance measure based on gradient information, referred to as Attention-weighted importance (see the “Gradient-based Importance and Attention-weighted Importance” subsection in Methods). Specifically, we compared Attention-weighted importance with the firing rate of input signals during training. The results showed that while Attention-weighted importance exhibited a significant positive correlation with the query-side training data, it demonstrated a significant negative correlation with the key-side training data. Furthermore, generated data showed no significant correlation with either the query or key sides (Figure 5e,f).

However, in Attention-weighted importance, the Key axis (columns) primarily holds information related to high-firing regions of the input signals, and the attention weights are determined based on how the Query side references this information. On the other hand, the Query axis (rows) is influenced by importance-based weighting, and the distribution of attention is readjusted due to the effect of the loss function. Previously, the model focused primarily on high-firing regions, but it has been adjusted to also pay attention to low-firing regions (see Methods).

In fact, modifying the loss function to the Dice function significantly improved the model’s predictive performance (Figure 3e,f). Within Attention-weighted importance, the Query side prioritized referencing distinctive information from high-firing regions of the input signals during training, primarily utilizing current and immediately preceding information. However, due to the influence of the Dice function’s evaluation, the Query side’s tendency in referencing Key-side information changed, and it was adjusted to also direct attention to distinctive features in past low-firing regions. As a result, the interaction between the priority derived from the input signals during training and the adjustment by the loss function contributed to the improvement in the model’s predictive performance.

Finally, we investigated the extent to which prediction performance deteriorated when the input data used for prediction was shuffled across different cells within the same time window (Figure 5g, right two bars). As a result, when shuffling was applied across cells, the model’s performance gradually declined as the window size expanded further into the past. However, even when 95% of the data was shuffled, the model retained a significant level of predictive accuracy (Figure 5g, rightmost bar). These results demonstrate that the model can still make reliable predictions even when the input data available for prediction is very limited.

## 4. Discussion

Here, we discuss the key findings of this study, categorized into technical advancements in methodology and neuroscientific insights.

### 4.1. Technical Advancements: The Role of Loss Function and Transformer Model

In this study, we adopted the Dice function as the loss for our Transformer model. While Transformer architectures and training toolkits have become ubiquitous, achieving high-precision generation of extremely sparse, binary spike data demands more than model choice alone. What appears to be a modest technical adjustment—the pairing of a single-layer Transformer with Dice Loss—proved beneficial in our experiments. To our knowledge, few studies have systematically evaluated this combination for spike-train generation; here we provide an empirical evaluation.

To our knowledge, this is the first study to combine these two elements specifically for the generation of neuronal spike trains. Despite the simplicity of the idea, no prior work recognized the unique synergy between the Transformer architecture and Dice Loss in addressing the extreme class imbalance inherent to neural spike data. As shown in Figure 3e,f, this combination dramatically enhances predictive performance, outperforming the specific baselines we tested (weighted cross-entropy and focal loss) on these datasets (Figure 3e,f).

It enables not only accurate within-modality generation (in vitro→in vitro, in vivo→in vivo), but also robust, bidirectional cross-domain generation between in vitro and in vivo conditions.

To understand the mechanisms underlying this high-accuracy generation, we first analyzed the core component of the Transformer model—the Attention Map. Our analysis revealed that the diagonal components of the Attention Map contributed significantly to model performance and that the query-side axis of the Attention Map exhibited a positive correlation with the firing rate of the input signals during training. To address a comprehensive interpretation of the model’s behavior beyond Attention Map, we also introduced a gradient-based importance measure, weighting the Attention Map with importance scores, referred to as Attention-weighted importance. A comparison between Attention-weighted importance and the firing rate of input signals revealed that while the query axis showed a significant positive correlation, the key axis exhibited a significant negative correlation.

Importance strongly reflects gradient information and is highly sensitive to the choice of loss function. This suggests that the performance improvement associated with loss function selection primarily influences the region between the input layer and the self-attention mechanism in the Transformer model. This adjustment enables the model to incorporate long-term historical information, facilitating more refined learning rather than relying solely on firing rates.

While the results suggested a limited role for the Attention Map in predictive accuracy, it is important to acknowledge its potential contributions to training stability and learning speed.

### 4.2. Neuroscientific Insights: Distinctive Brain Regions

Despite achieving high-accuracy predictions and generation, certain data exhibited particularly noteworthy characteristics.

The first notable finding concerns the meaningful multi-region mapping shown in Figure 4f. This mapping clearly captures the expected spatial relationships among data points in multiple aspects. For instance, the two data points measured from the secondary motor cortex are closely aligned and surrounded by regions associated with the motor cortex both of in vitro or in vivo data. Additionally, the in vivo and in vitro data are spatially separated into two distinct clusters on the left and right. These semantically meaningful embeddings represent the relative relationships between data points, suggesting how activity transitions from one data point to another. In this study, we demonstrate cross-generation from brief spontaneous activity in both traditional in vitro data and the in vivo data provided by the International Brain Laboratory. Choosing the optimal embedding dimensionality is always a challenging problem. If one spatially maps neural-activity similarity by directly comparing it to actual spatial distances, both in vitro and in vivo points would naturally lie in three dimensional space. Hence, embedding the two datasets (in vitro and in vivo) in three to four dimensions is reasonably justified for their comparison in this work. However, should the number of datasets under comparison grow to three, four, five, or more, new justification will be required to determine whether a three dimensional visualization remains appropriate.

The second region of interest is the lateral pre-optic area (LPO) in vivo. This region appeared to be among the stronger seeds for predicting other brain regions in our dataset. Please note that the abbreviation “LPOR” used in Figure 4 and Table 1 refers to the left postrhinal area, which is a different region. The abbreviation for the lateral preoptic area in this context is “LLatPreopt”.

The LPO, a hypothalamic nucleus, is one of the most extensively connected subdivisions of the hypothalamus: in the mouse it projects to and receives input from over 200 gray-matter regions, with intra-hypothalamic connections being especially prominent (Hahn et al., 2022). For example, it sends output to the lateral habenula (LHb), septal nuclei, ventral tegmental area (VTA), dorsal raphe nucleus, and the rostromedial tegmental nucleus (RMTg) (Gordon-Fennell et al., 2020). It also receives input from the lateral septal nucleus (LS), which itself is a cortical hub region receiving hippocampal projections. Moreover, the LPO has reciprocal connections with the subiculum as well as CA1 and CA3 fields of the hippocampus.

The LPO is involved in both reward processing and the regulation of sleep–wake states (Saper et al., 2005; González et al., 2016). Since sleep and wakefulness are fundamental behavioral states that entail wide-ranging shifts in brain function, the LPO’s connectivity to arousal centers is critical. Recent work has also shown functional coupling between the LPO and the reward system: stimulation of the LPO suppresses GABAergic neurons in the VTA while increasing the firing rate of dopaminergic neurons (Gordon-Fennell et al., 2020). Intriguingly, the LPO participates in an integrative circuit linking the nucleus accumbens—the ventral striatum involved in “positive” reward valuation—to the lateral habenula, which encodes aversive reward prediction errors.

Taken together, the LPO receives cognitive and sensory inputs from cortex and hippocampus and, via its outputs to brainstem and forebrain arousal and reward centers, functions as a hub that regulates global brain-state variables such as sleep–wake cycles, motivational drive, and homeostasis. This extensive functional connectivity allows LPO activity to “mirror” patterns of activity across diverse regions and, when necessary, influence network-wide dynamics. In other words, by virtue of its broad inputs, the LPO may internally represent spontaneous cortical activity patterns in miniature and emit potent control signals tightly linked to cortical dynamics. It is noteworthy that our data-driven analysis highlighted the LPO as a region of interest.

### 4.3. Future Challenges: Expanding the Range of Applications

Based on these findings, two efficient strategies can be proposed.

First, the “proximity map” based on relative similarities between datasets expressed as a network diagram in Figure 4f is very important. This is because the proximity map encodes relative relationships among datasets, when a measurement is sparse or unavailable the framework can propose concrete substitutes—for example, selecting nearby datasets, mixing closely related datasets, or using the generative model to translate between conditions along the map’s geometry. These are approximations of missing conditions rather than de novo creation, and they require task-specific validation.

From the standpoint of the 3Rs, this capability is a foundational technology that leads to the reduction of redundant experiments when existing results have been independently reproduced, while also prioritizing follow-up studies. That said, decisions to forgo new experiments should be based on predefined criteria and ethical review, and this method can be regarded as a technology that provides evidence for deliberations in that ethical review. In the future, as the number of nodes (datasets) in the network increases and network density grows within the informatics framework, the accuracy of generating non-existent data will steadily improve.

Second, prioritizing LPO measurements before expanding to other brain regions may enhance predictive accuracy and experimental efficiency because the Lateral preoptic area data provides good seeds for generating neural activity across many regions. This strategy aligns with the 3R principle (Replacement, Reduction, Refinement) in animal research and could improve efficiency in human neurophysiological studies. The complete explanation of why this region’s neural activity can serve as such a versatile learning data seed (independent of the aforementioned proximity) remains unclear. As our understanding deepens, generation without requiring target data is expected to become increasingly feasible.

In the future, when considering the contribution of the LPO, some researchers in the life sciences may envision experiments involving optogenetic stimulation of this region. However, what is truly essential is to elucidate the “codes” utilized in the process of generating neural activity. In other words, it is crucial to uncover the information acquired by artificial neural networks through learning. To deepen our understanding of such phenomena, we expanded the interpretational scope of the Transformer’s internal structure from attention maps to attention-weighted importance. Future challenges include further expanding this analysis and extracting and conducting detailed analysis of features from attention maps and Importance that contribute to prediction-generation.

In addition, it is important to improve methods for enhanced accuracy. Simple improvements include adding Position encoding to the Transformer model. As the computational method itself is scalable, expanding computational resources, such as computer memory, to increase the analyzable number of cells and time duration is also an important direction. While we performed mutual generation based on spontaneous activity, extending this to generate in vivo brain activity during stimulus presentation is another crucial direction. This can naturally be pursued by inputting in vivo spontaneous activity and stimulus information into a multi-modal AI model.

## 5. Conclusions

In conclusion, this study demonstrates a practical, high-precision method for generating neural activity and introduces a novel framework for dynamic integration and comparison of existing experimental datasets. By mapping spike-train dynamics rather than static connectivity, our approach may help reduce redundant replications of past experiments (“Reduction” of the 3Rs) and is intended to complement—rather than replaces—new experimental work. We will continue to improve accuracy, extend prediction horizons, and deepen our mechanistic understanding. In the future, this work may provide a basis for comparing analyses across animal and human datasets, but further validation will be required.

## Supplementary Materials

Not applicable.

## Author Contributions

MS was responsible for the study design, data-sharing negotiations, data analysis, and manuscript preparation.

## Institutional Review Board Statement

Ethical approval. All animal procedures complied with institutional and national regulations. For in-vitro experiments, the use of C57BL/6J mice (3–5 weeks old) was approved by the Kyoto University Animal Experimentation Committee and performed in accordance with Kyoto University guidelines. A total of six mice were used in this study. The total number of mice used for the in vitro experiments in this study is six. For the in-vivo datasets analyzed here, all procedures were conducted in accordance with local laws and with approvals from the Animal Welfare Ethical Review Body of University College London; the Institutional Animal Care and Use Committees of Cold Spring Harbor Laboratory, Princeton University, the University of California at Los Angeles, and the University of California at Berkeley; the University Animal Welfare Committee of New York University; and the Portuguese Veterinary General Board, as written in International Brain Laboratory et al. 2023, 2025. The total number of mice used for the in vivo experiments is also six.

## Informed Consent Statement

Not applicable.

## Data Availability Statement

All original and generated datasets used in this study, together with related figures, are publicly available at Mendeley Data (“GenerativeNeurosci_TFDice”, V1; doi: 10.17632/kf65cvmtbz.1). The analysis and generation code is available at https://github.com/ShimonoMLab/GenerativeNeurosci_ML-TrDic. For third-party source datasets, please refer to the original repositories and citations listed in the Methods and References.

## Abbreviations

ACSF: artificial cerebrospinal fluid
AUC: area under the curve
IBL: International Brain Laboratory
LPO: lateral preoptic area
ROC: receiver operating characteristic.

## Acknowledgments

MS is deeply grateful to Gaelle Chapuis of the International Brain Laboratory (IBL) and to Dora Angelaki, Tom Mrsic-Flogel, and Sonja Hofer for generously sharing invaluable neuronal activity data recorded with Neuropixels probes.

